# Genetic improvement of wheat early vigor promote weed-competitiveness under Mediterranean climate

**DOI:** 10.1101/2020.09.27.315531

**Authors:** Shlomi Aharon, Aaviya Fadida-Myers, Kamal Nashef, Roi Ben-David, Ran N. Lati, Zvi Peleg

## Abstract

Chemical weed-control is the most effective practice for wheat, however, rapid evolution of herbicide-resistant weeds threat food-security and calls for integration of non-chemical practices. We hypothesis that integration of GAR dwarfing-genes into elite wheat cultivars can promote early vigor and weed-competitiveness under Mediterranean climate. We develop near-isogenic lines of bread wheat cultivars with GAR dwarfing genes and evaluate them for early vigor and weed-competitiveness under various environmental and management conditions to identify promising NIL for weed-competitiveness and grain yield. While all three NILs, OC1 (*Rht8,12*), ZC4 (*Rht12,13*), and BNMF12 (*Rht12*), responded to gibberellic acid, they exhibited differences in early vigor. Greenhouse and field evaluation highlighted OC1 as a promising line, with significant advantage in early vigor over its parental. To facilitate accurate and continuous early vigor data collection, we applied non-destructive image-based phenotyping approaches which offers non-expensive and end-user friendly solution for selection. NIL OC1 was tested under different weed density level, infestation waves, and temperatures and highlight the complex genotypic × environmental × management interactions. Our findings demonstrate the potential of genetic modification of dwarfing genes as promising approach to improve weed-competitiveness, and serve as basis for future breeding efforts to support sustainable wheat production under semi-arid Mediterranean climate.

## 1. Introduction

Wheat (*Triticum* spp.) is one of the most important grain-crops in the world, with annual production of 734 million tons across 214 million hectares worldwide (https://faostat.fao.org). Weeds are recognized as the main biotic factor limiting grain production, causing globally over 34% of crop yield losses [1]. This negative effect on wheat productivity acting either directly (i.e., costs of management such as labor, equipment and chemicals) or indirectly (i.e., via competition on resources, reduction in grain yield and quality, increasing the processing cost, and as secondary host for pets). While herbicides are the most cost-effective and efficient practice for weed control in wheat, in recent years, it has become less efficient due to the rapid evolution of herbicide-resistant weeds [2], and climate change depended tolerance [3]. The threat posed by herbicide-resistance on food security calls for a new paradigm of weed control in large-scale crop production agro-systems [4]. Thus, there is an urgent need to integrate non-chemical cultural weed control practices, such as mechanical cultivation, flaming, higher sowing density, modification in inter- and intra-row spacing, intra-row cover crops, and no-till [5-7].

A cost-effective strategy to improve weeds control in wheat could be achieved by breeding new characteristics that promote crop to outcompete and suppress weeds (i.e. enhance competitive ability). Crop competitiveness against weeds can be divided into: *suppression*, the ability of cultivar to reduce the fitness of a competitor weed, or *tolerance*, the ability of the crop to tolerate competition from weeds and maintain high yields [8]. Higher genetic variation for weed suppression as compared with weed tolerance has been reported. Moreover, weed tolerance showed strong G × E interaction (i.e. low heritability), suggesting greater opportunity for the selection of improved weed suppression in wheat [9]. Weed-competitiveness includes complex traits such as early vigor, leaf area index (LAI), number of tillers, canopy architecture (erect/ spread) and canopy height. All these traits were found correlated with weed suppression of cereals [5, 10-12]. However, tradeoffs between weed competition and yield potential can also affect the breeding selection of these traits.

Early crop vigor (i.e., canopy area produced early in the growing season) is considered crucial trait for semi-arid Mediterranean wheat agro-system by promoting optimal field stand, faster establishment and canopy closure and increased radiation-use efficiency (RUE) [13, 14]. Moreover, it decrease soil water evaporation from soil surface, which can support higher grain yields under Mediterranean conditions. Early seedling vigor is also an important component in crop ability to compete with neighboring plants [15-18]. Plant characteristics associated with high early vigor and weed competitiveness includes grain or embryo size [19, 20], leaf characteristics (width, length and area), seedling emergence rate [14], rapid early growth rate [21], optimization of tillers number [17], large specific leaf area index during vegetative growth [14, 22] root architecture [23-25].

The modern semi-dwarf wheat cultivars post the “Green Revolution” are characterized by higher grain yield and improved harvest index [26, 27]. However, the deployment of gibberellin (GA) insensitive (GAI) dwarfing genes, *Reduce height* (*Rht*)-*B1b/Rht-D1b*, resulted in decrease cell size, shorter coleoptile, and reduce leaf size during early growth stages, and as consequence poor weed competitiveness. This low competition ability, among other factors, makes chemical herbicides the prime method to control weeds in wheat. Genetic improvement of early vigor might be achieved via several traits such as increased grain or embryo size, wider leaf and longer coleoptiles. One strategy to counter this, is using alternative GA-responsive (GAR) dwarfing genes, which reduce plant height while avoiding the reduction effect on early vigor and coleoptile length, and suggested a means to improve early vigor [28-30].

To date, only a few breeding programs focused on improving early vigor (i.e., crop competitiveness) to promote sustainable integrated weed management (IWM) practice [5, 6, 31]. Increased interest in breeding wheat with improved weed suppression is growing in response to the evolution and rapid expansion of herbicide resistant weed populations, the needs of organic agro-systems, and the limited access of smallholder farmers in the developing countries to chemicals [32]. Our working hypothesis is that integrating alternative dwarfing GAR genes into modern bread wheat cultivars can improve early vigor and weed competitiveness and promote grain yield in Mediterranean agro-system. The objectives of the current study were to (***i***) develop near isogenic line of elite bread wheat cultivars with alternative semi-dwarf genes (***ii***) test whether GAR introduction can contribute to wheat early vigor and weed competitiveness, and (***iii***) establish field-based evaluation of promising NIL for weed competitiveness and grain yield.

## 2. Material and Methods

### 2.1 Plant material

Three bread wheat cultivars (Omer, Zahir and Bar-Nir) were crossed with donor parents harboring alternative dwarfing genes (Chuan-Mai 18 [*Rht8*] and Marfed Dwarf [*Rht5, Rht13*]; Table S1). The F1 plants were grown to produce F2 seeds. The F2 seedlings were selected based on DNA markers for the desired genes (Table S2) and backcrossed with the recurrent parent for three generations, followed by six selfing to produce BC3F6 (Fig. S1). In each generation of crossing, the lines were phenotypically selected based on their response to exogenous application of gibberellic acid as described previously [28]. At the last stage, the developed near isogenic lines (NILs) were genetically characterized to validate their genetic composition. The NILs genetic composition is as follow: OC1, *Rht8* and *Rht12*, BNMF12, *Rht12* and ZC4 still have the *Rht*-*B1b* (GAI) as well as two alternative GAI genes *Rht12* and *Rht13* (Table S3).

### 2.2 Image-based quantification of early vigor among the NILs in control conditions

Uniform seeds of three NILs (OC1, ZC4 and BNMF12) as well as their recurrent parental lines (*cv*. Omer, Zahir and Bar-Nir, respectively) were sown in 1 liter pots, filled with clay soil (57% clay, 23% silt, and 20% sand, on a dry-weight basis and 2% organic matter). Pots were placed in controlled growth chamber with photosynthetic light (temperature 16°C) in complete randomized design, with five replicates. The experiment was conducted at the Newe Ya’ar Research Center. Plants were watered by an automated mini-sprinkler irrigation system (VibroNet(tm) 25 L/H, Netafim, Isreal) as needed, and fertilized every two weeks with N:P:K (20:20:20).

Each plant was phenotyped for the projected shoot area (PSA) from 10 days after sowing (DAS) until 20 DAS (every other day, total five evaluations). Plants were imaged using a Nikon D7200 digital single-lens reflex camera (DSLR) fitted with a Nikkor 18-140 mm lens (Nikon Corporation, Tokyo, Japan) that was placed 90° relative to the ground (Fig. S2A). The PSA of the plants were automatically extracted as described before [33]. The shoot-related pixels were classified from their surrounding background (Fig. S2B) and their exact area was evaluated using absolute area units (cm^2^). At the end of the experiment (30 DAS), the above ground material was harvested, oven-dried at 70°C for 72 h and weighted to obtain shoot dry weight (DW).

### 2.3 Evaluation of NILs panel for early vigor in the field

Field experiment was carried out to validate the improved early vigor of promising NILs compared to their recurrent parents at the experimental farm of the Hebrew University of Jerusalem in Rehovot, Israel (34°47′ N, 31°54′ E; 54 m above sea level). The soil at this location is brown-red, degrading sandy loam (Rhodoxeralf) composed of 76% sand, 8% silt and 16% clay. The experiment was held in a paired sample complete block design, with seven replicates. Seeds of three NILs, (OC1, ZC4 and BNMF12), as well as their recurrent parental lines (*cv*. Omer, Zahir and Bar-Nir, respectively) we sown using plot seeder)Wintersteiger AG), with12 m long plots, at seed density of 200 plants m^2^.

Wheat early vigor was estimated using the spectral parameters Excess Green Vegetation Index (ExG) [33] that were extracted via image-data acquired from Unmanned Aerial Vehicle (UAV). Flights conducted 30 m above ground level, resulting in ground sampling resolution of ∼0.6 cm/pixel, using a DJI Phantom 4 Pro UAV (Shenzhen, Guangdong, China). The UAV is equipped with 20 M Pixel RGB camera with wavelengths at 620-750 nm (R, red), 495-570 nm (G, green) and 450-495 (B, blue). Mission planning was conducted using DJI Ground Station Pro (GSP) software. Flight were with 80% image overlap along flight corridors. Then, Pix4DMapperPro desktop software (Pix4D SA, Switzerland, http://pix4d.com) was used to generate ortho-mosaics. Six ground control points (GCP) geolocated with Real Time Kinematic (RTK) survey precision were used to georeference the ortho-mosaics. The Pix4D processing options were done according Pix4D’s “3D Maps” template version 4.1.10. The ExG values were derived from the georeferenced ortho-mosaic GeoTIFFs that were generated from the UAV flights using the following equation:

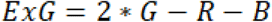

where R represents the return on the red channel, G represents the return on the green channel and B represents the return on the blue channel. Plot-level ExG means from UAV’s were created in ArcGIS^®^ 10.6 (ESRI, Redlands, USA). Shape files with an area of 6 m^2^ containing annotated single plot polygons were generated. ExG plot means were generated using the Zonal Statistics as table function in ArcGIS.

### 2.4 Characterization of weed-competition ability

This experiment aimed to compare the weed-competition ability between NILs. Uniformed seed were sown in 20 liter (38 × 32 × 16 cm) plastic containers, filled with a mixture of fine peat and sand (ratio 1 : 4, respectively), at a density equivalent to of 250 plants m^−2^. Seed were sown in three rows, with eight seeds per row (in total 24 seeds) simultaneously to weed density treatment. Due to relatively low germination rate of weeds, we used cultivated ryegrass (*Lolium multiflorum*) and black mustard (*Brassica nigra*) as model weeds (germination rate >95%). Four ryegrass and four mustard seeds (total eight seeds per container) were used to mimic weed density of 70 weeds m^2^. The experiment conducted using a complete randomization design, with six replicates for each combination: weed-free wheat, wheat and weed competition and only weeds. Containers were placed in the net-house and plants were grown for six weeks. At the final stage of the experiment, wheat and weeds shoots were harvested, oven-dried at 70°C for 72h and weighed to obtain shoot DW.

### 2.5 Characterization of wheat and weeds competiveness under various environmental and management conditions

This experiment aimed to test the effect of various environmental and management conditions, late timing of weed infestation, weed-densities and temperature, on the weed competiveness ability of selected NILs. Experiments were held in net-house as described above, in a complete randomization design, with six replicates. Late weed infestation treatment was applied by sowing the weed seeds three weeks after wheat sowing. Weed-density treatments were applied by sowing weeds at two rates equivalent to 70 and 140 weeds m^−2^, 8 and 16 seeds of ryegrass mustard per container, respectively. Temperature treatments were applied by dividing the containers between two net-houses with average of 16°C and 20°C. The later net-house had plastic roof that induced higher air temperature. Temperatures were recorded throughout the trial using HOBO data logger.

### 2.6 Evaluation of selected NILs for weed competitiveness in the field

Field experiment was carried out to validate the advantage of specific NILs, OC1, in term of weed competitiveness over its recurrent parent Omer. Experiment was held at the Newe-Ya’ar research station (32.706069’N, 35.183073’E) using randomized block design with six replicates. Wheat seeds were sown at a density of 250 plants m^−2^, i.e., ten seed rows, with 500 seeds per plot (2 m^2^). Weed infestation was based on the natural seed bank. Weed-free and wheat-free treatments were used as control plots. The weed-free plots were manually weeded weakly. Three and five weeks after germination, weed shoots were harvested from randomly selected 625 cm^2^ sample to obtain shoot DW. Then, the PSA of wheat in the same sample areas (after weeds removal) was evaluated to determine the response of each genotype to weed competition. The PSA data was extracted using image data as described above by DSLR camera. Here, images were taken paralleled to the soil surface and the PSA was evaluated for the sampled 625 cm^2^ area. Additionally, ExG values of the hand weeded control plots of Omer and NIL OC1 were evaluated using UAV as described in section 2.3. Evaluations were held two, three and four weeks after germination (WAG). At the end the experiment, grain yield and 1000 kernel weight (TKW) were calculate to each plot. Grains were extracted using a laboratory threshing machine (350-LD, Wintersteiger) and weighted.

### 2.7 Statistical analyses

All statistical analyses were conducted using the JMP^®^ ver. 15 statistical package (SAS Institute, Cary, NC, USA). To test the differences between each NIL and its recurrent parent we used a t test (*P*≤0.05). The first field experiment (Rehovot) was analyzed as paired sample t-test blocks design. The rest of mean comparison between NIL pairs was done using t-test.

## 3. Results

### 3.1 Introgression of alternative dwarfing genes affect wheat early vigor dynamics

To test the effect of the introgressed GAR dwarfing genes on plant-growth rate dynamics during early stages, we characterize the set of NILs in comparison with their recurrent parent under controlled conditions. In general, most NILs showed improved early vigor as expressed by PSA compared with their parental line (Fig. 1A-C). NIL OC1 showed the highest improvement, with significantly higher PSA value compared to Omer early as 12 DAS (3.3 *vs*. 2.0 cm^2^, respectively). This advantage of NIL OC1 increased throughout the experiment, with PSA value reaching 12 cm^2^, two fold higher compared to Omer 20 DAS (Fig. 1A). NIL ZC4 showed also a significant advantage over its recurrent parent Zahir in PSA values throughout the experiment, starting 12 DAS, and up to 9.5 cm^2^ compared to 5.5 cm^2^, respectively at 20 DAS (Fig. 1B). NIL BNMF12 showed similar performance as its parental line Bar-Nir through the experiment (Fig. 1C). To further support the image-based phenotyping, shoot DW was analyzed 30 DAS. Both NILs OC1 and ZC4 accumulated significantly higher biomass values compared to their parental lines (median values of 80 *vs*. 125 mg and 65 *vs* 110 mg, respectively), whereas NIL BNMF12 had similar values (80 mg) (Fig. 1D-F).

**Figure 1.**
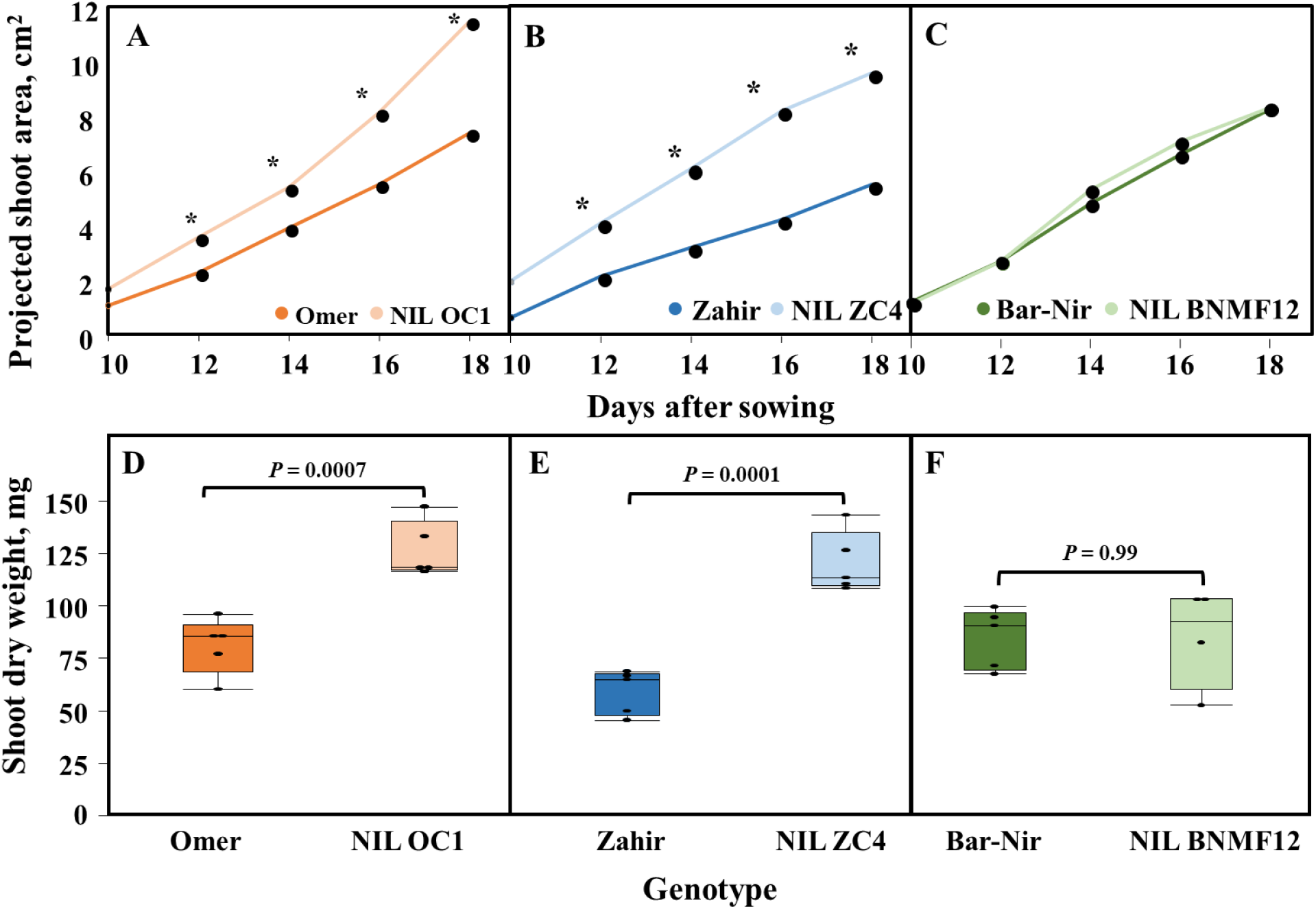
Longitudinal projected shoot area of (**A**) Omer and NIL OC1, (**B**) Zahir and NIL ZC4, and (**C**) Bar-Nir and NIL BNMF12. * indicate significant difference between cultivar and its NIL using t-test (*P*≤0.05). Comparison between shoot dry weights evaluated 30 days after sowing of (**D**) Omer and NIL OC1, (**E**) Zahir and NIL ZC4, and (**F**) Bar-Nir and NIL BNMF12.

In order to validate the performance of the selected NILs, we conducted field experiment. The differences in the ExG values between each NIL and its parent were analyzed, and as Figure 2 shows, NIL OC1 was the only line that showed significant (*P*=0.045) improvement under the uncontrolled field conditions. Furthermore, the difference median value was −5 (ranging between – 17 to 5) with only one replicate with <0 value, suggesting the increase of NIL OC1 early growth compared to Omer is prominent (Fig. 2). NILs ZC4 and BNMF12 showed significantly lower early vigor ability (*P*=0.009 and *P*=0.01, respectively) compared to their parents with median values >8 for both lines.

**Figure 2.**
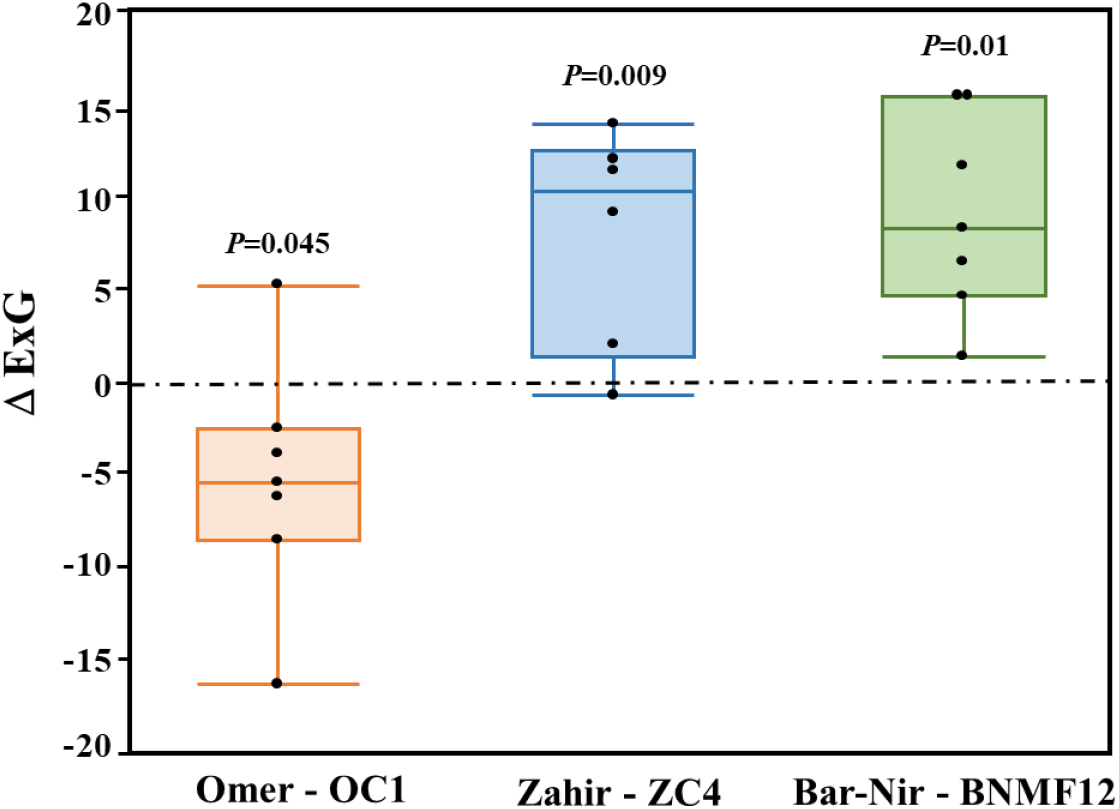
Paired sample t-test comparison of the ExG vegetation index differences (Δ) between the three set of cultivars and their respective NIL evaluated in the Rehovot field experiment.

### 3.2 Early vigor promote higher weed competitiveness

To test if the genetically improved early vigor of the NIL also contributes to their weed-competitiveness ability we conducted competition experiment by applying weed-infested conditions. The weeds competitiveness ability was evaluated by two parameters: the wheat shoot DW accumulation and the detection of any inhibition in weeds development and productivity (i.e. biomass DW). In general, the NIL OC1 showed improved competitiveness ability as compared with its recurrent parent Omer (42 DAS; Fig. 3). Under competition with weeds, OC1 had higher canopy density as compared to Omer (Fig. 3A), and improved suppression ability of both weed species, ryegrass (Fig. 3B) and black mustard (Fig. 3C). These weeds growth was stunted and had slower phenological development compared to weeds that were grown with *cv*. Omer.

**Figure 3.**
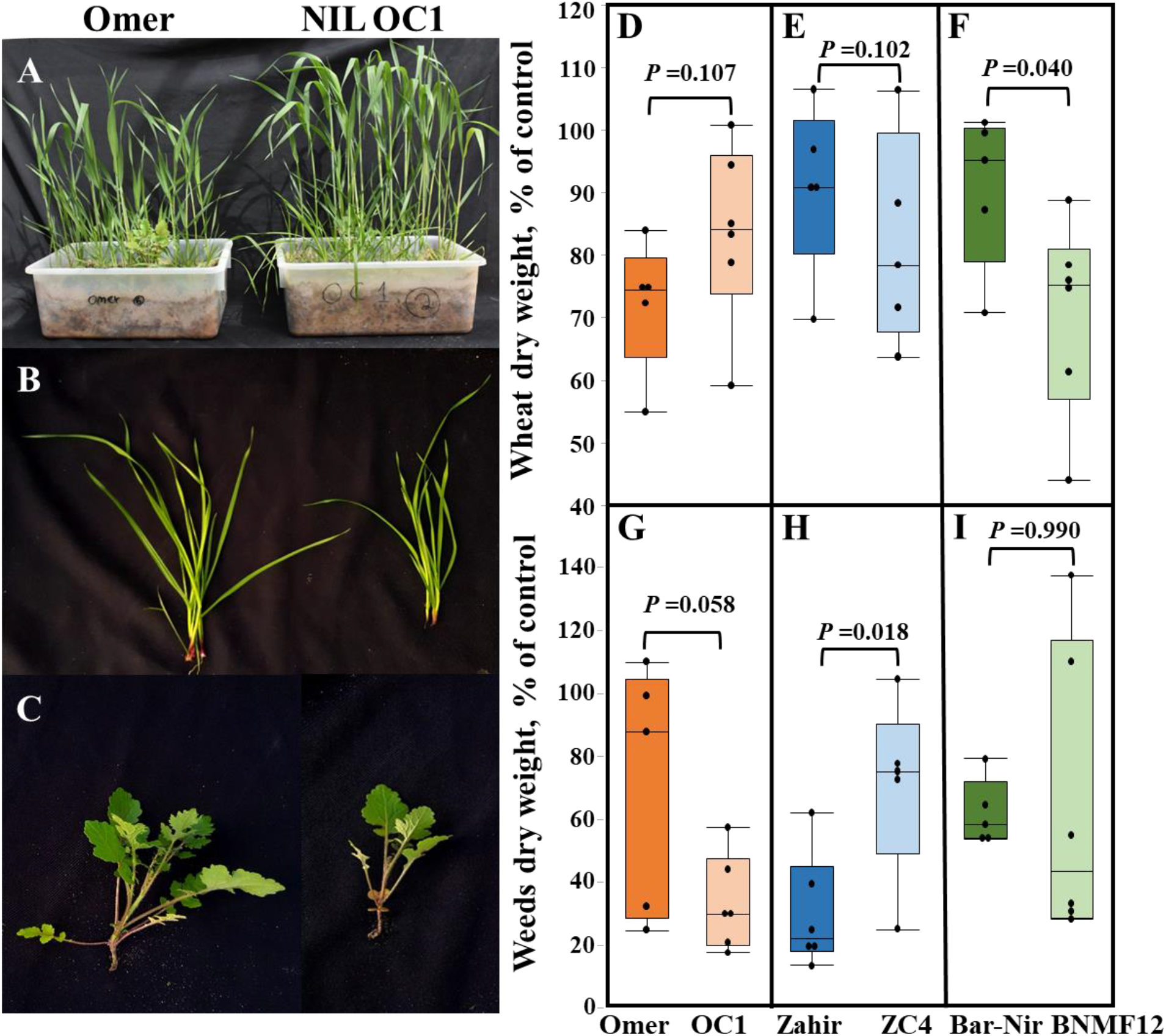
Effect of competition with weeds on wheat development. (**A**) A representative photo of wheat plants grown under completion with weeds. Weeds phenology under competition with wheat (**B**) ryegrass (*Lolium multiflorum*) and (**C**) black mustard (*Brassica nigra*). Wheat biomass under competition with weed relative to performance under weed-free conditions for (**D**) Omer and NIL OC1, (**E**) Zahir and NIL ZC4, and (**F**) Bar-Nir and NIL BNMF12. Weeds biomass under competition with wheat relative to only weed for (**G**) Omer and NIL OC1, (**H**) Zahir and NIL ZC4, and (**I**) Bar-Nir and NIL BNMF12.

Quantitative analysis of wheat shoot DW at the end of the experiment (six WAS) showed similar advantage of NIL OC1, however, with no significant difference compared to Omer (Fig. 3D). The other NILs ZC4 and BNMF12 were more sensitive to weed interference compared with their recurrent parents and exhibited lower shoot DW (Fig. 3E-F). The inhibition of weeds development in a response to the wheat competition of NIL OC1 was the strongest, resulting in significant lower weed DW (*P*=0.058) at OC1 plots compared to plots of Omer (Fig. 3G). The two NILs that were more sensitive to the weed infestation, BNMF12 and ZC4, had differential effect on weed development. NIL ZC4 was a weak competitor and the weeds DW that was sampled from its plots was significantly higher (*P*=0.0178) compared to weeds sampled in Zahir plots (Fig. 3H). Weeds DW at NIL BNMF12 and Bar-Nir plots did not differ significantly (*P*=0.99) (Fig. 3I).

### 3.3 Strong interaction between genotype and management or environmental factors affect plants early vigor

Based on the improved early vigor and weed-competitiveness we selected NIL OC1 for detailed characterization. The stability and robustness of this improved early vigor and weed-competition ability of OC1 was tested under various environmental conditions (temperature and weed density) that mimic the Mediterranean basin and managements (timing of weed infestation) (Fig. 4). Since many field in the Mediterranean agro-system exhibited patchiness in term of weed infestation density, we tested the ability of newly developed NILs under two density levels (70 and 140 weeds m^2^). While under low weed density NIL OC1 showed similar shoot DW compared to Omer, under the high weed density level, OC1 DM was higher than Omer (*P*=0.0421). Under both weed densities, NIL OC1 exhibited improved competition ability, which is expressed in lower weed DW at NIL OC1 compared to Omer, low (*P*<0.0583) and high (*P*<0.0535) densities (Fig. 4A-B).

**Figure 4.**
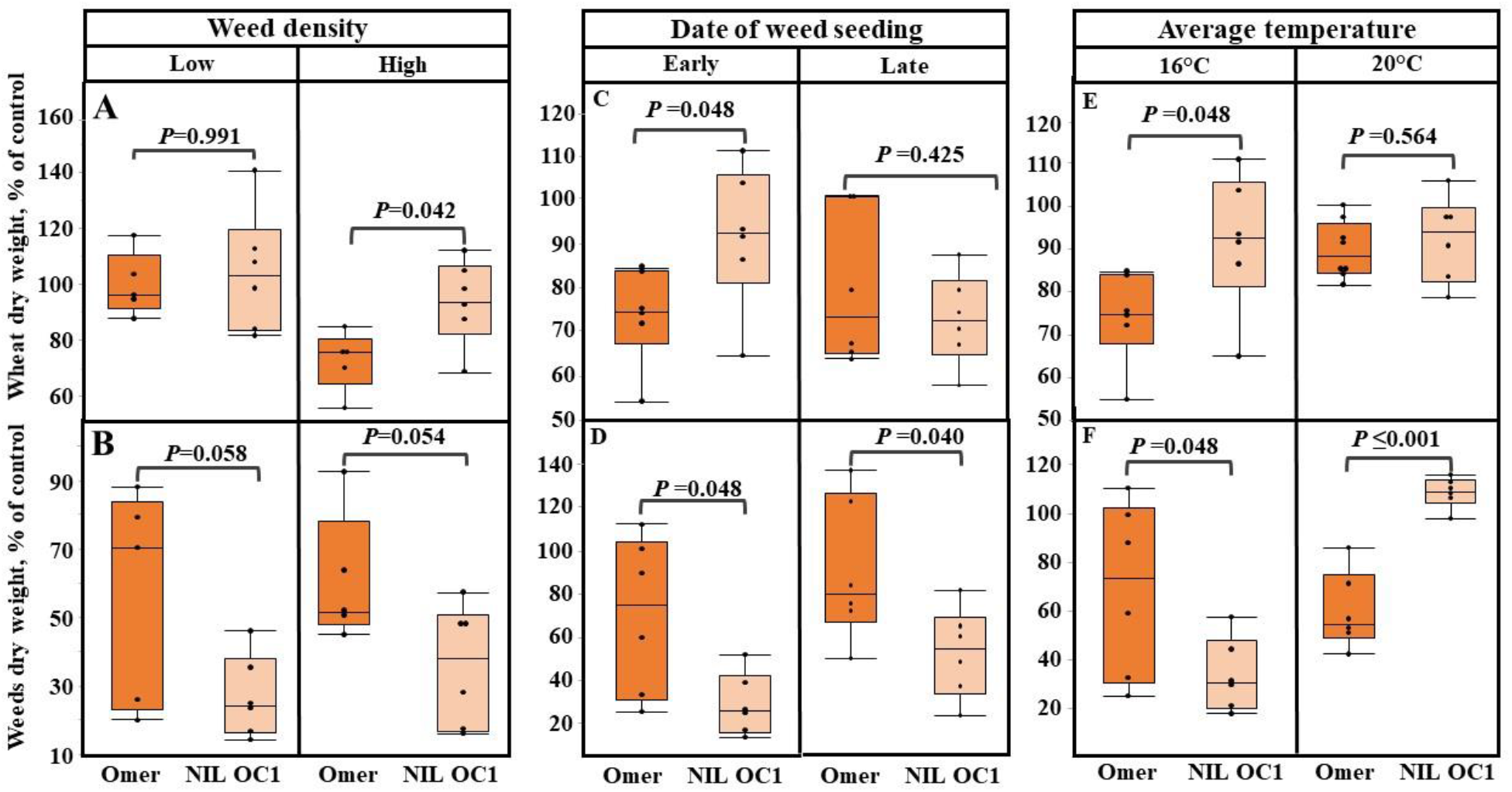
Effect of environmental and management conditions on competitiveness of *cv*. Omer and NIL OC1. Relative biomass of (**A**) wheat and (**B**) weeds under low and high weed density. Relative biomass of (**C**) wheat and (**D**) weeds under early and late weed seeding timing. Relative biomass of (**E**) wheat and (**F**) weeds under low (16°C) and high (20°C) temperatures. Comparison between Omer and OC1 conducted by t-test (*P*≤0.05).

Weed management in the rain-fed wheat agro-system in the Mediterranean basin include usually a first herbicide application at crops 3-4 leaves stage (to control weed population established pre-crop emergence). A second application will take place later in the season in order to control the secondary flush of weed germination. To mimic this secondary germination flush after herbicide application, we applied post-crop emergence infestation of weeds (when crop is at 3^rd^ leaf stage(. In term of wheat shoot DW, NIL OC1 showed significant advantage over Omer only under early weed infestation. In both cased, however, the NIL showed improved weed-competition ability, as expressed in significant reduction in weed DW (23 *vs*. 77% and 55 *vs*. 80% of weed DW compared to control for early and late weed infestation, respectively; Fig. 4C-D).

Temperature can significantly impacts wheat plants development, and consequently, the weed-competition ability. In recent years, climate change increased the intensity of extreme temperature event during the winter, and therefore, we tested the performance of both lines under normal (16°C) and heat wave (20°C) conditions. Under the lower temperature (16°C), OC1 shoot DW was higher than Omer (*P*=0.047) (Fig. 4E). Consequently, DW of weeds seeded in OC1 plots were lower compared to weeds seeded in Omer plots (*P*=0.0483; Fig. 4F). Under the higher temperature (20°C) this trend was inversed. No significant differences (*P*=0.564) were observed between NIL OC1 and Omer shoot DW. The weeds dry weights that were seeded in NIL OC1 plots were significantly higher than weeds that were seeded in Omer plots (*P*<0.0001). No inhibition was observed for weeds that were seeded in NIL OC1 plots with median value of 100% of control compared to 50% inhibition for weeds that were seeded in Omer plots (Fig. 4F).

### 3.4 NIL OC1 exhibited higher early vigor and weed-competitiveness under field conditions

To validate the improved early vigor and weed-competition of NIL OC1 over Omer we conducted field experiment under real field conditions. In general, the NIL OC1 exhibited advantage over Omer in both vegetative growth (i.e. early vigor) parameters and in weed inhibition. Three weeks after germination the OC1 plants were more developed and had denser canopy compared to Omer, and this advantage increased with time. The advantage of OC1 was also expressed in lower levels of weed developed in the OC1 plots as compared to Omer (Fig. 5A). This observation was further supported by quantitative analysis. Two WAG, OC1 showed only a slight trend of advantage over Omer in shoot DW (both had median ∼50% of control) and weed competition ability (median of 40% and 80% of control, respectively), which was not significant (Fig. 5B-C). Later on in the season, five WAG, the improved competition ability of OC1 was more evident and it was reflected via both parameters. Wheat PSA of OC1 was significantly (*P*=0.01) higher compared to Omer, with median values of 75% and 55% of control, respectively. DW of weeds that developed in OC1 plots were significantly (*P*=0.05) lower than weeds that developed in Omer plots, with median values of 20% and 50% of control, respectively (Fig. 5B-C). In parallel, analysis of vegetation index (ExG) with UAV imaging support the same pattern (Fig. S4).

**Figure 5.**
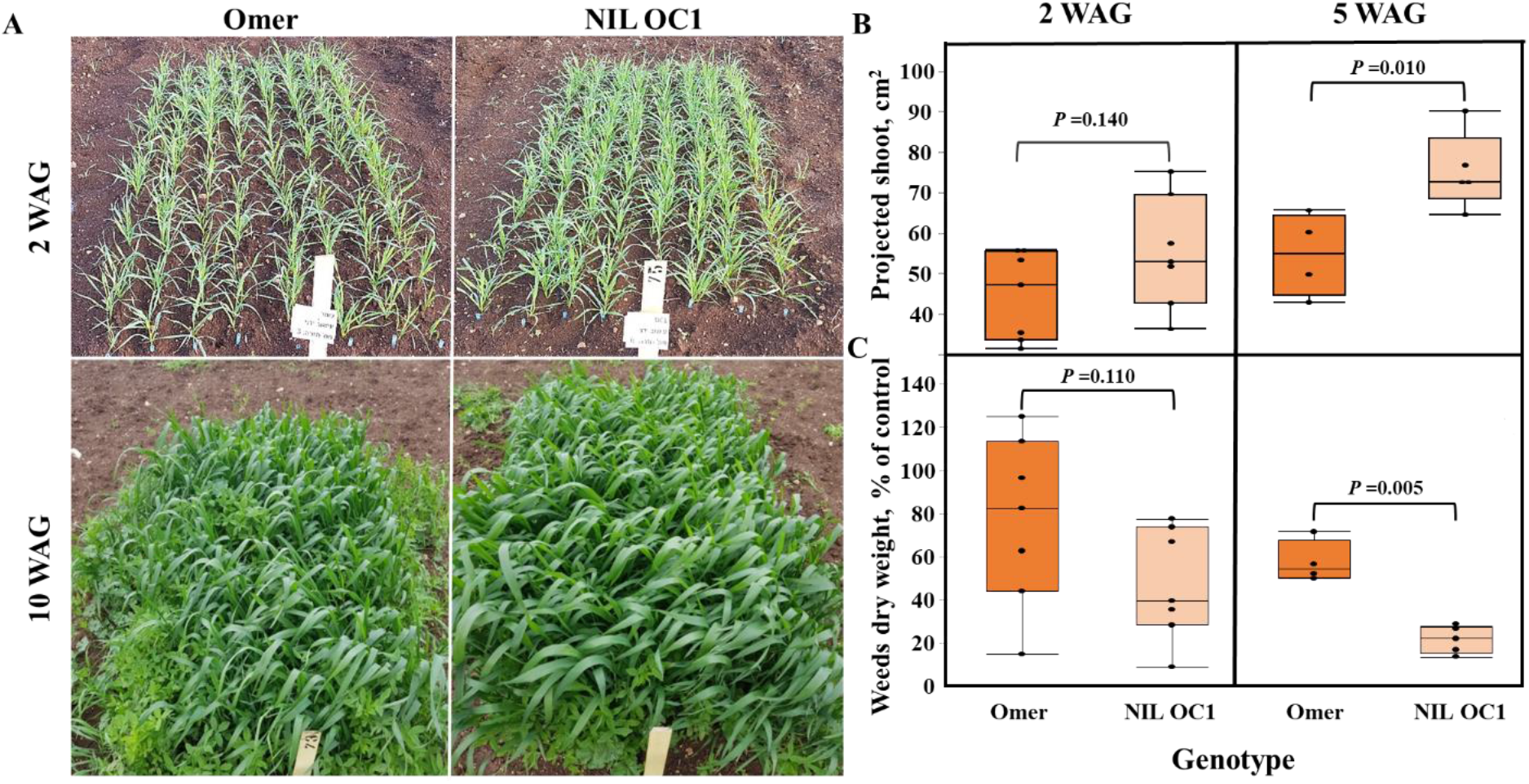
Field validation of early vigor advantage in weed-competitiveness. (**A**) A representative photo of Omer and NIL OC1 in the field 2 (up) and 10 (bottom) weeks after germination (WAG) under field conditions. (**B**) Projected shoot area of Omer and NIL OC1 2 (left) and 5 (right) WAG. (**C**) Relative weed dry weight under competition with wheat Omer and NIL OC1, 2 and 5 WAG. Data was evaluated in the Newe Yaar field experiment. Comparison between Omer and NIL OC1 conducted by t-test (*P*≤0.05).

In general, grain yield and TKW of NIL OC1 didn’t show difference from its recurrent parent Omer (Fig. 6). This comparison was held under three weed competition scenarios: full competition (non-treated control), no competition (hand weeded control) and partial competition (herbicide treatment). For example, the manual weeding treatment resulted with grain yield of ∼0.52 kg m^−2^ for both NIL OC1 and Omer, and with respective grain weight of 43 and 39 g, nonetheless, with lack of significance trend (*P*=0.309).

**Figure 6.**
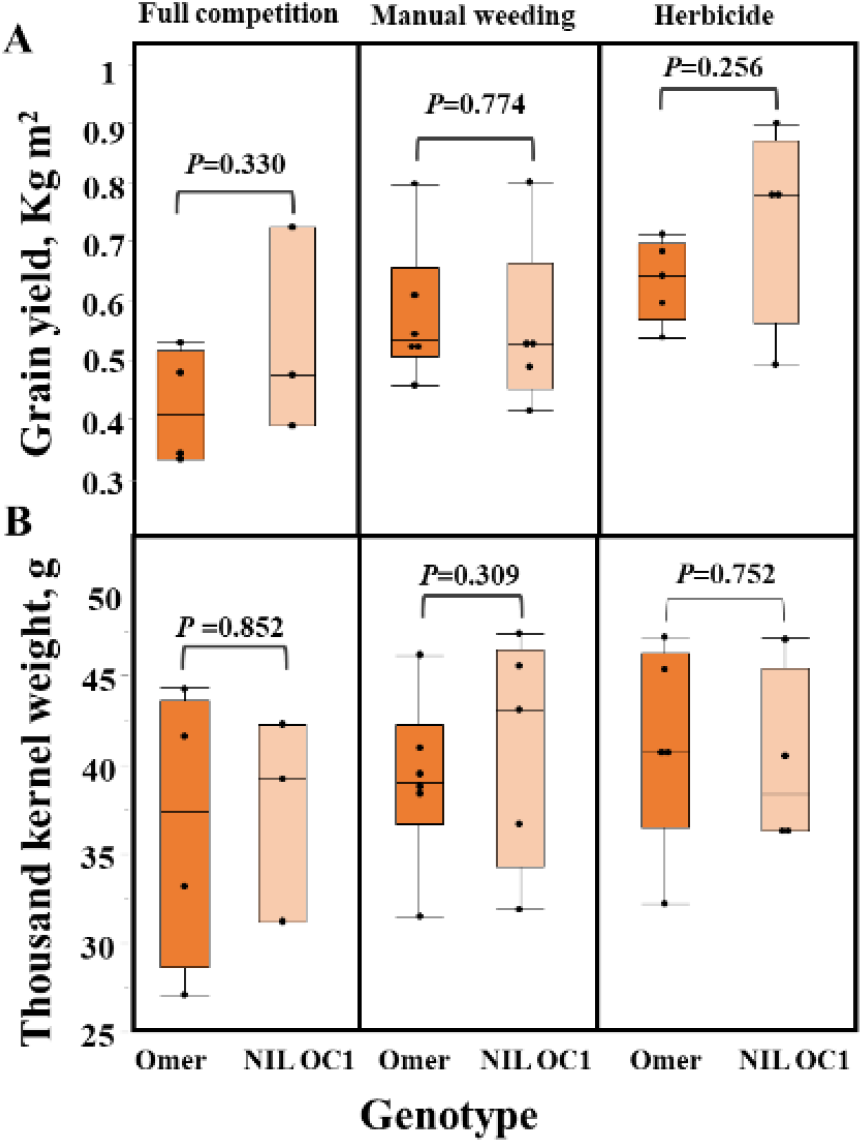
Effect of early vigor and weed-competitiveness on wheat yield. (**A**) Grain yield and (**B**) thousand kernel weight of Omer and NIL OC1 under competition with natural weeds in the field, manual weeding and herbicide application. Comparison between Omer and OC1 under each treatment conducted by t-test (*P*≤0.05).

## 4. Discussion

Effective weed control will play a key role in leveling the yield gap associated with future climate change in the Mediterranean basin. While herbicide is the most efficient strategy for weed control, this by no mean provides hermetic solution to the complex challenge weeds provides for farmers [34]. Moreover, in recent years herbicide-resistant weed populations are evolving rapidly as a consequence of increasing selection pressure imposed by modern agricultural management practice [2]. Thus, there is an urgent need to develop alternative and integrated weed management practices. Here we demonstrate the potential of genetic modification of plant architecture which enables the crop to better compete with weeds. In our view this is an economically and environmental-friendly solution that could reduce or even eliminate the need for chemical or mechanical weed control [35].

Our current study introduce several alternative GA-responsive dwarfing genes into background of elite bread wheat cultivars adapted to semi-arid Mediterranean climate. While all three NILs are responsive to GA, they showed various growth pattern. NIL OC1 (*Rht8, 12*) showed significant advantage in growth over the recurrent parent Omer under controlled greenhouse and field conditions. NIL ZC4 which still poses the *Rht-B1b* dwarfing gene with two alternative genes (*Rht12, 13*) showed significant advantage in growth over the recurrent parent Zahir under controlled greenhouse conditions, however it was not expressed under field conditions. NIL BNMF12 (*Rht12*) didn’t show any difference from its recurrent parent Bar-Nir, under either controlled greenhouse or field conditions (Figs. 1-2). In previous study’s *Rht12* was found to confer strong dwarfism (45%) and significant delay in heading [29]. As *Rht12* is shared among all the tested GAR NILs in this study we can conclude that in Mediterranean spring wheat background those prominent phenotypic effects is hardly expressed with all GAR NILs showing similar phenology and heights (OC1 was indeed the highest NIL, but was only 20% higher than Omer). The non-reducing early vigor effect of the individual *Rht* GAR genes involved in this study (*Rht8, 12, 13*) is well documented [29, 36, 37]. However, data on the combine effect on early vigor of two GAR alleles in a specific genetic background is sparser. Our results suggest that *Rht12* by itself results in a similar reduced early vigor as *Rht1* (NIL BNMF12). When combined with *Rht13* early vigor in the NIL is recovered at list under controlled conditions. Combining *Rht12 and Rht8* results in full early vigor recovery both in pots and on filed. A mechanism of enhanced cell files across the leaf blade contributed to wider leaves and greater early vigor of *Rht*8-carring lines [38, 39] and might also explain the high weed competitiveness of OC1 found in this study.

Breeding of plant architecture traits is long, tedious and labor-intensive task, and involved in many cases collection of destructive parameters (i.e. plant biomass). As consequence, it hinder capturing the plant longitudinal nature dynamics which can provide better phenotypic resolution required for adequate data interpretation [40]. To facilitate effective, accurate and continuous data collection, we applied a non-destructive image-based phenotyping approaches. We integrated proximal (consumer grade camera) and remote (UAV) evaluation strategies to uncover the plant growth dynamic, as reflected by PSA. Both strategies were able to distinguish, with high confidentiality, between the genotypes longitudinal growth dynamics (Fig. S2). Moreover, the system sensitivity enables to detect differences in growth dynamics already after 12 days after sowing (Fig. 1). Thus, these tools offers high-throughput, non-expensive and end-user friendly phenotyping solution for research and breeding of early vigor traits that is relevant to small scale laboratory experiments as well as large scale field-based phenotyping of germplasm.

Weed competitiveness is complex trait expressing the dynamic interaction between crop-plant and weeds. This dynamic is rooted not only in direct factors (i.e. crop-plant *vs*. weeds) but also via indirect factor, either environmental (i.e. temperature, precipitations) or management-related (i.e. field stand and timing of chemical application). From an IPM perspective, weed density level is an outcome of soil seed bank, on-field management history and environment. Different weed infestation waves (highly dependent on herbicides practices) will affect the developmental stage mostly affected by wheat-weed competition. Interestingly, NIL OC1 exhibited improved weed competitiveness, as expressed in weeds DW, under both weeds densities (high and low) and timing (pre- and post-crop emergence). However under the controlled conditions NIL OC1-enhanced early vigor (shoot DW) was detected only under high weed density or under pre-emergence weed competition (during emergence). In the first case, NIL OC1 enhance early vigor may be driven by density dependent below ground crop-weed allele-chemical signals supported by recent reports by Kong et al. [41]. Wheat competiveness varietal scores is reported to maintain ranking stability across different weed densities ([17] and citation therein) although this was concluded in context of high-weed densities. Lower weed density (70 weeds m^2^) were chosen for this study under which NIL OC1 shoot DW was not higher then control. Under field conditions, NIL OC1 shoot DW superiority detected along all developmental stages (Fig. 5), in agreement with previously reported data [42].

In a simulation study of field data, [43] it was found that wheat early vigor and specific leaf area (SLA, a component of early vigor) affect crop performance and yield with increased impact under high temperatures. In the current study NIL OC1 showed advantage in early vigor only under low temperatures (typically for the Mediterranean basin), in contrast with low shoot DW and higher weed DW in higher temperature (Fig. 4). Similar trends were observed under control environment, with high temperatures (24°C compare to 8°C) resulting in reduce wheat shoot growth [44]. These findings further emphasis the important of extending the experimental context (both environmental and agronomical) and the need to capture accurately G×E response. Notably, in previous study, increased air and soil temperature affected negatively coleoptile length in both GAI and GAR genetic backgrounds [42, 45, 46]. As coleoptile length is an important component of early vigor [GAR genotypes (e.g., NIL OC1) exhibiting enhanced coleoptile length compared to GAI wheat lines (e.g., Omer)] this negative effect of temperature might also contribute to the establishment and consequently to low shoot DW of NIL OC1 under high temperatures. Pre-breeding effort targeted for high temperature scenarios should concentrate on wide screening for wheat genotypes with enhanced weed competitiveness less responsive for heat as was previously suggested for long coleoptile trait [45]. This can also apply for the future climate change in the Mediterranean climate and projected increased temperatures.

While we were able to provide evidence that NIL OC1 have clear advantage in early vigor under control conditions, to validate the improved early vigor and weed-competition we conducted field experiment under real field conditions. The performance of NIL OC1 outcome Omer in both vegetative growth (i.e. early vigor) and weed inhibition (Figs. 5, S4). However, NIL OC1 did not exhibited any advantage in grain yield and quality traits under the various weed competition scenarios [full competition (non-treated control), no competition (hand weeded control) and partial competition (herbicide treatment)] (Fig. 6). The development of NIIL OC1 involved replacing the GAI with GAR dwarfing genes. It was suggested that in case GAR dwarfing effect is more subtle possible off set in HI optimization might occur potentially reducing grain yield (e.g., *Rht13* or *Rht8,13* combination) [30]. Here we show that while NIL OC1 is indeed taller its improved weed-competitiveness was not associated with any yield penalty [42]. Moreover, the development of wheat ideotype with improved weed-competitiveness can support IWM less dependent on herbicides. Yet, farther research to test the interaction of NIL OC1 with different environmental and management practices is needed.

## 5. Conclusions and future perspective

Modifications of wheat morphology (i.e. the ideotype concept; [47]) is essential to increase wheat weed-competitiveness and productivity under the projected climate change. The improvement of early vigor can be achieved by various complimentary strategies either implementing genetics or advanced managements. Here we demonstrate the potential of genetic modification of GAR dwarfing-genes as promising new approach to improve weed-competitiveness. Moreover, wheat with higher crop early vigor is likely to have greater water-use efficiency in environments like the semi-arid Mediterranean basin, where a large component of total water use is soil evaporation [48]. Overall, our findings expand the knowledge of the genetic basis underlying the G×E×M interaction associated with early vigor and weed-competitiveness. Thus, integration of early vigor trait into wheat breeding program for semi-arid condition can support more sustainable (i.e. reduce chemical use) and more efficient resources use (reduce water demand) to promote stable crop production.

## Acknowledgments

This study was supported by the Chief Scientist of the Israeli Ministry of Agriculture (grants # 12-01-0005 and 837-0134-13). We thank members of the Peleg, Ben-David and Lati labs for their invaluable assistance in the field experiments.

