## Supplementary material for "Genetic improvement of wheat early vigor promote weed-competitiveness under Mediterranean climate": SI data

**Table S1.** List of modern bread wheat (recurrent parents) and donor parents.

| | Genotype / cultivar | Species | Origin |
|---|---|---|---|
| **Wheat landraces (donor GAR *Rht* genes)** | | | |
| | **Chuan Mai-18** | Bread Wheat | China |
| | **Mara** | Bread Wheat | Italy |
| **Modern wheat cultivars** | | | |
| | **Omer** | Bread Wheat | Israel |
| | **Zahir** | Bread Wheat | Israel |
| | **Bar-Nir** | Bread Wheat | Israel |

**Table S2.** List of genetic markers used for marker assisted selection.

| *Rht* gene | Chromosome | Primer | Sequance | Size |
|---|---|---|---|---|
| **GA-Insensetive dwarfing genes** | | | | |
| *Rht-B1b* | 4BS | BF-MR1 | GGTAGGGAGGCGAGAGGCGAG<br>CATCCCCATGGCCATCTCGAGCTA | 237bp |
| *Rht-D1b* | 4DS | DF-MR2 | CGCGCAATTATTGGCCAGAGATAG<br>CCCCATGGCCATCTCGAGCTGCTA | 254bp |
| **Alternative GA-Responsive dwarfing genes** | | | | |
| *Rht5* | 3BS | BARC102 | GGAGAGGACCTGCTAAAATCGAAGACA<br>GCGTTTACGGATCAGTGTTGGAGA | 200bp |
| *Rht8* | 2DS | WMC503 | GCAATAGTTCCCGCAAGAAAAG<br>ATCAACTACCTCCAGATCCCGT | 275bp |
| *Rht9* | 5AL | BARC151 | TGAGGAAAATGTCTCTATAGCATCC<br>CGCATAAACACCTTCGCTCTTCCACTC | 220bp |
| *Rht12* | 5AL | WMC410 | GGACTTGAAAAGGAAGCTTGTGA<br>CATGGATGGCATGCAGTGT | 114bp |
| *Rht13* | 7BL | WMS577 | ATGGCATAATTTGGTGAAATTG<br>TGTTTCAAGCCCAACTTCTATT | 13bp |

**Table S3.** Table S3. List of near-isogenic lines (NILs), their recurrent and donor parental line and their composition of dwarfing genes.

| NIL | Recurent parent | Donor line | *Rht* genes |
|---|---|---|---|
| **OC1** | Omer | Chuan-Mai 18 | *Rht8*, *Rht12* |
| **ZC4** | Zahir | Chuan-Mai 19 | *Rht1*, *Rht12*, *Rht13* |
| **BNMF12** | Bar-Nir | Marfed Dwarf | *Rht12* |

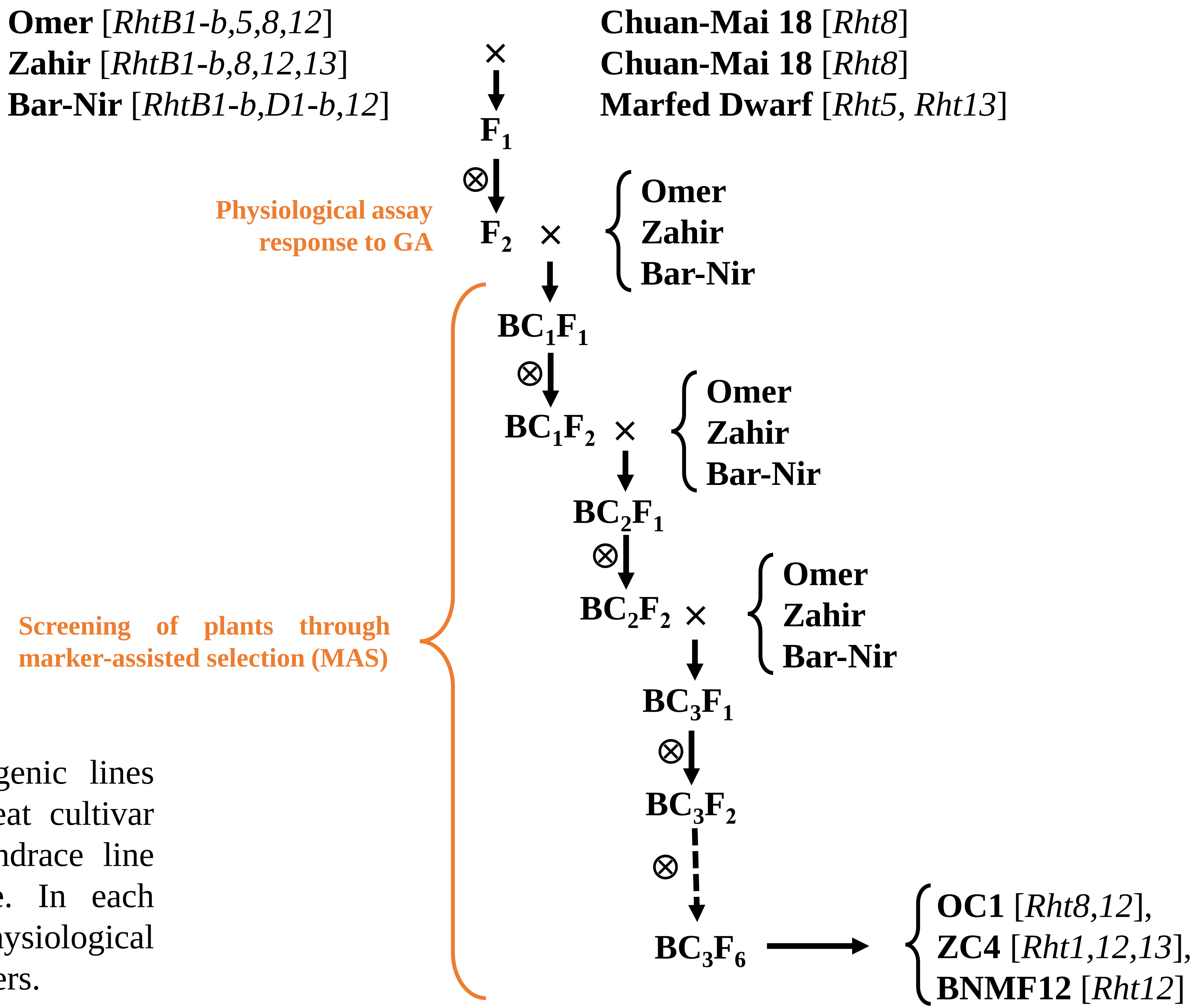


**Figure S1.** Development of near isogenic lines (NILs). Cross between elite bread wheat cultivar carry the GAI dwarfing genes and landrace line carry GAR alternative dwarfing gene. In each generation, plants were screened with physiological assay (response to GA) and genetic markers.

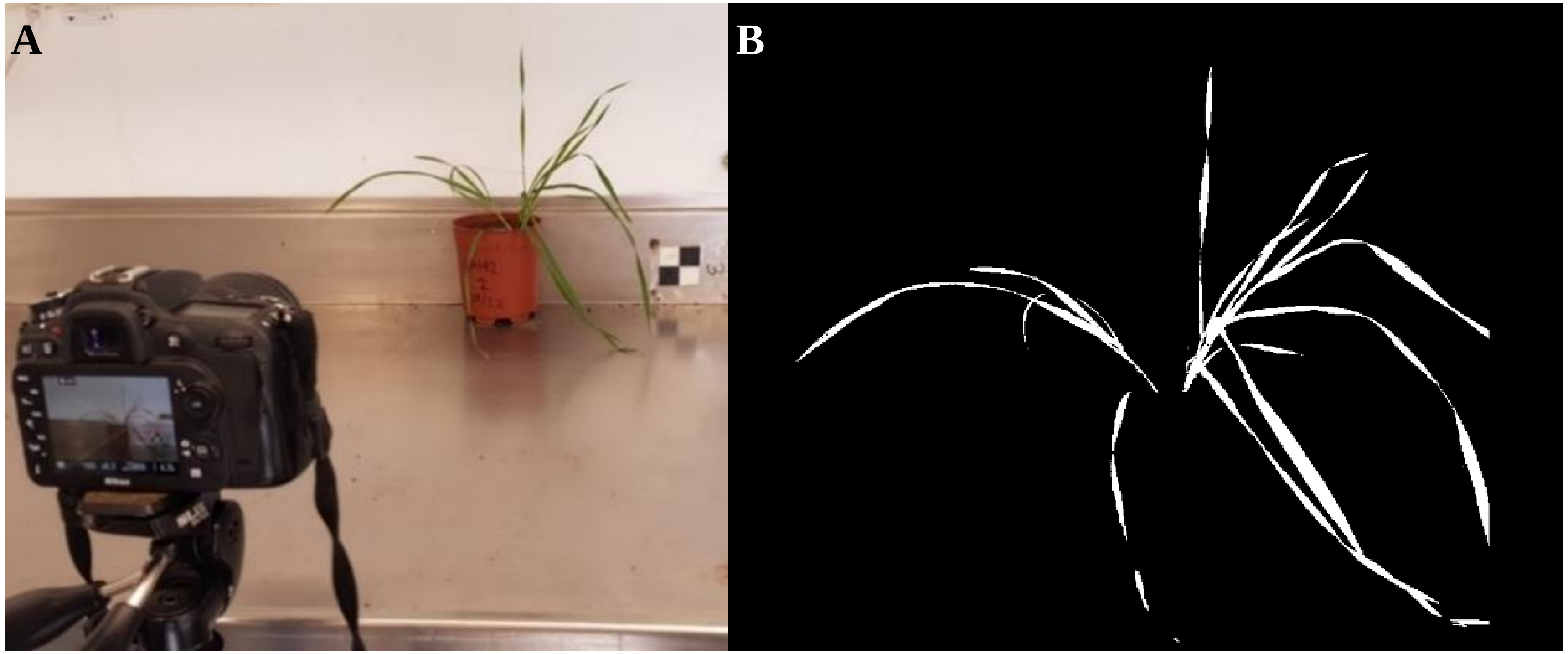


**Figure S2.** (**A**) Capturing single RGB images of individual plant. (**B**) assessment of projected shoot area (PSA).

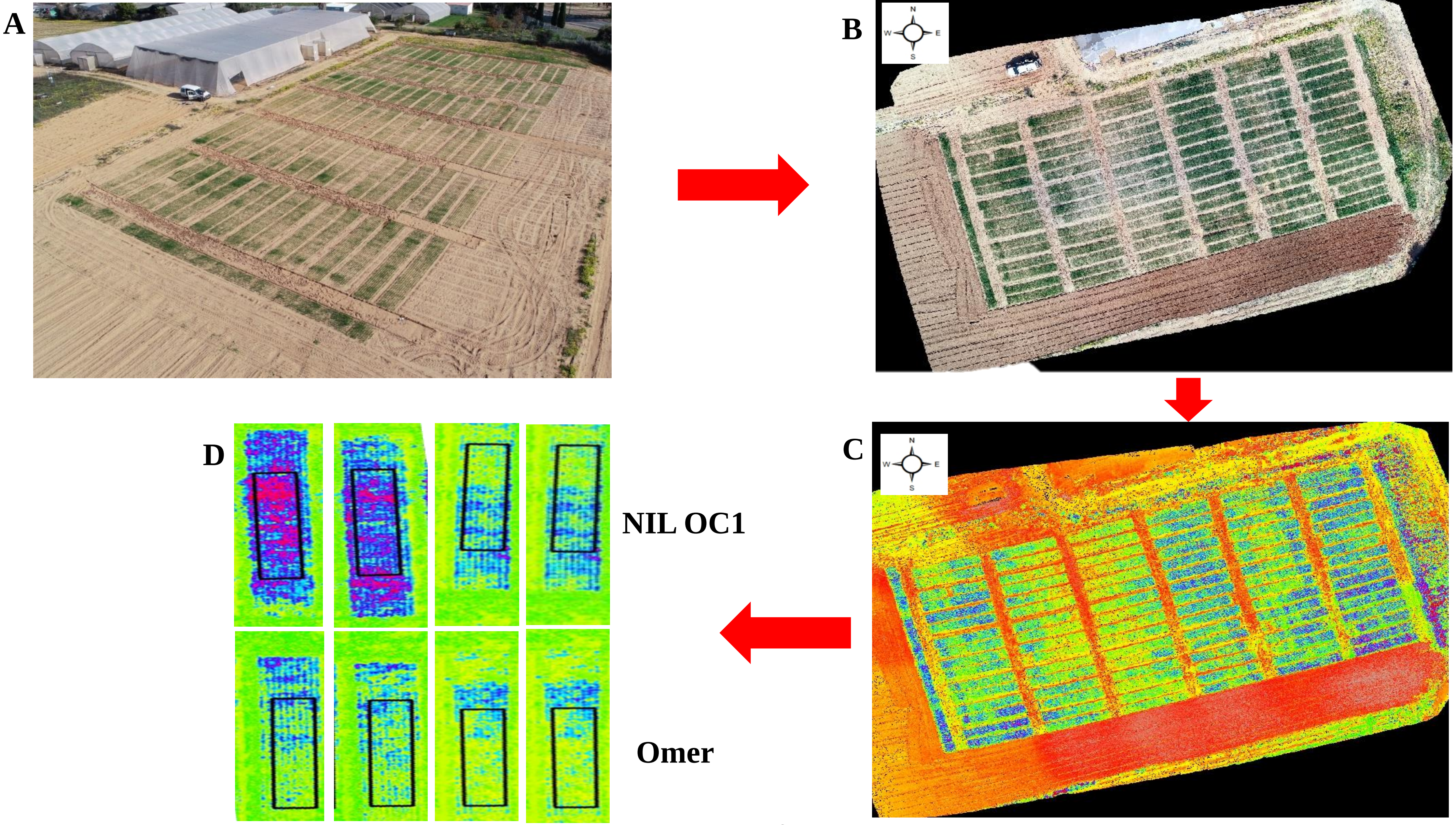


**Figure S3.** Excessive Green (ExG) evaluation pipeline in the field trials. (**A**) Imaging the experimental site using RGB camera mounted on UAV, following by (**B**) stitching the images into ortho-mosaic and (**C**) application of the ExG vegetation index. Finally, (**D**) placing the polygones on the exact plot area and extraction of the average ExG values.

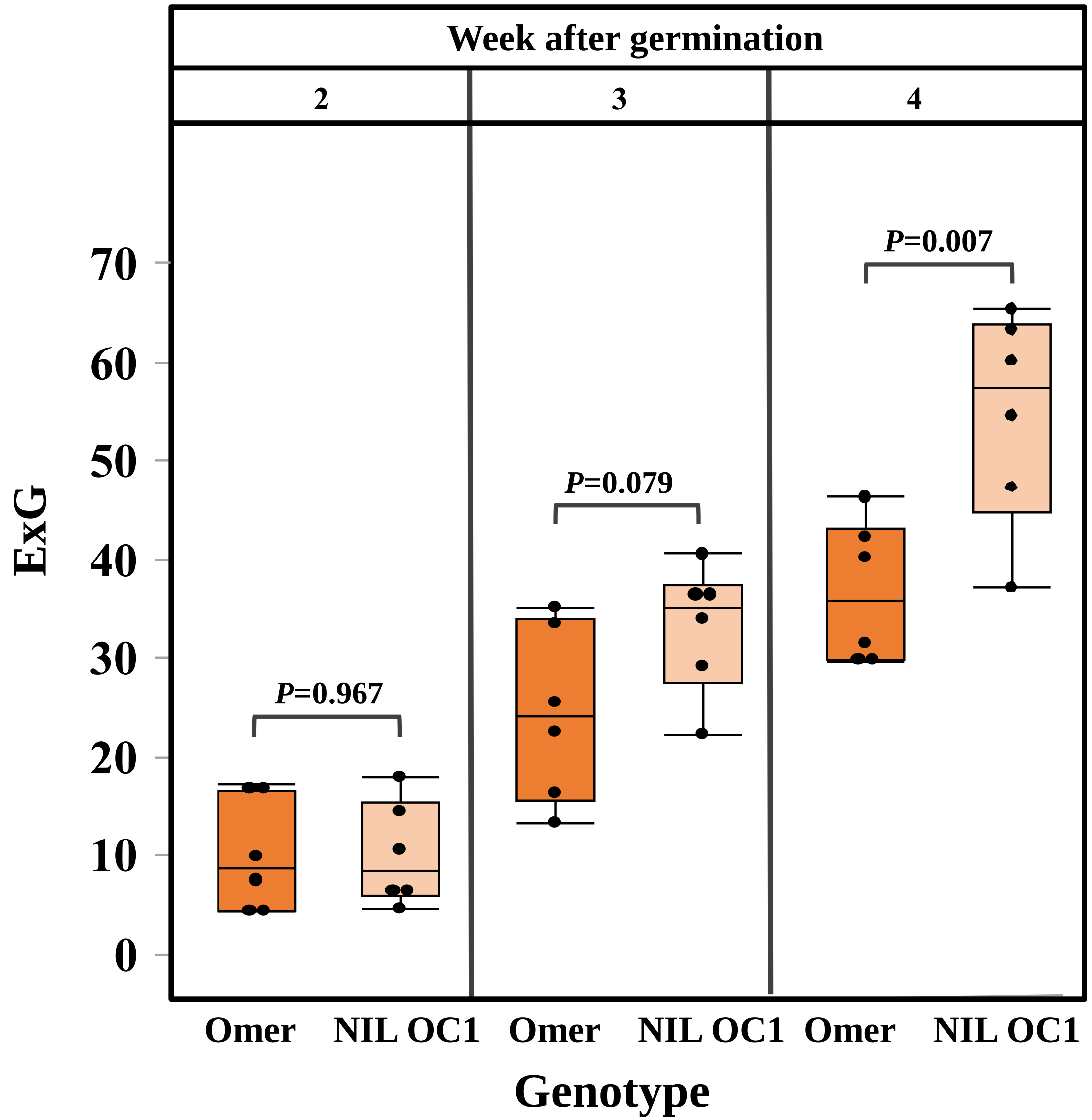


**Figure S4.** ExG vegetation index of *cv*. Omer and NIL OC1in the field along the developmental stages, from week 2 to week 4 after germination evaluated in the Newe Yaar field experiment.
